# Mitochondria at synapse utilize fatty acids as a bioenergetic fuel source

**DOI:** 10.1101/2025.05.26.656116

**Authors:** Bernardo Cetra Antunes, Andreia Faria-Pereira, Joana Gonçalves-Ribeiro, Sara Costa-Pinto, Sandra H. Vaz, Vanessa A. Morais

## Abstract

Mitochondria process glucose, glutamine, and fatty acids (FAs) to produce ATP, with fuel choice dependent on tissue-specific metabolism. The brain harbors two distinct mitochondrial populations—synaptic and non-synaptic. While glucose is the primary fuel for brain bioenergetics, the role of FAs remains elusive. A preliminary proteomic analysis revealed that synaptic mitochondria favor FA metabolism, corroborated by biochemical and respiratory assays showing their higher capacity for β-oxidation and greater respiratory flexibility compared to non-synaptic mitochondria. Additionally, synaptic mitochondria showed higher capacity for FA uptake and less susceptibility to inhibition of the carnitine shuttle system. In neurons, oxygen consumption rate assays indicate that medium to long-chain FA fueling enhances neuronal respiratory flexibility and ATP content, while whole-cell patch-clamp recordings show that long-chain FA fueling sustains increased pre-synaptic activity. Our findings demonstrate that FAs can contribute effectively to synaptic metabolism under normal physiological conditions, where there is a constant demand for energy.

## INTRODUCTION

Synapses host the most energy demanding processes in the brain [1], and under disease conditions, synaptic damage often precedes neuronal death [2]. Synapses, often located in considerable distances away from the cell body, in some cases measurable in meters, possess mitochondria with an unique fingerprint to facilitate and secure the homeostasis of synaptic transmission [1]. Synaptic mitochondria differ from remaining brain mitochondria, often termed non-synaptic, in lipidome and proteome [3]–[5], enzymatic expression and activity [6]–[8], as well as in shape and size [1]. Indeed, the fission machinery and smaller size of synaptic mitochondria are crucial for axonal colonization and homeostasis [9]. Moreover, what these specific differences between sub-compartmentalized mitochondria dictate remains to be clarified.

One claim, that is withstanding, is the fact that maintenance of neuronal homeostasis entails a constant energetic supply to support neurotransmission, axonal and dendritic transport, synthesis of neurotransmitters, and protein synthesis, being the dependency on glucose well documented [10]. However, if glucose alone is enough to keep neuronal circuitry functioning properly, or if additional fuel resources are required remains to be clarified. Fuels such as glutamate [11] and fatty acids [12] have been shown to sustain neuronal metabolism to a certain extent. The relevance of fatty acid oxidation (FAO) in the brain has been undermined due to decreased enzymatic efficiency of FAO enzymes in brain mitochondria when compared to mitochondria from other tissues [10], [13]. On the other hand, metabolism of fatty acids induced metabolic coupling between neurons and astrocytes [12], [14]. With an energetic output that surpasses glucose on more than 2-fold [10], FAO may be important for neurons, even if only in extreme conditions.

The question remains whether glucose alone is sufficient for synaptic mitochondria to cope with the stressful and energy-demanding conditions characteristic of the neuronal periphery, or if additional fuel sources, such as fatty acids, are necessary to maintain synaptic homeostasis. To understand the importance of FAO in neurons, particularly at the synapse, we assessed firstly the metabolic capacity of synaptic mitochondria towards this fuel, in comparison to non-synaptic mitochondria, via enzymatic, oxygen consumption and substrate usability assays, and secondly the effect of FAO in the biology of mouse primary neurons, from respiration to electrical activity. Our data reveals that synaptic mitochondria have a higher propensity for FAO when compared with non-synaptic mitochondria, demonstrating increased enzymatic efficiency, metabolism, uptake capacity, and respiratory output upon stimulation of the electron transport chain. At the cellular level, fueling with fatty acids enhanced neuronal respiration and overall ATP levels, ultimately sustaining increased presynaptic activity, as evidenced by increase spontaneous postsynaptic currents (sPSCs).

## RESULTS

### The proteome of synaptic mitochondria shows upregulation of proteins involved in the FAO

A proteomic screen previously performed in the host laboratory comparing synaptic and non-synaptic mitochondria isolated from C57BL/6 mice (Supplementary Fig.1) (also described in [15]) revealed increased levels of FAO proteins in synaptic mitochondria (Fig. 1A and B). These proteins are involved both in the carnitine shuttle that imports long-chain fatty acids to mitochondria (carnitine-palmitoyl transferase 2 - CPT2), as well as in the four steps of β-oxidation (acyl-CoA dehydrogenase family member 9 - Acad9, enoyl-CoA hydratase, short chain 1 -Echs1, 3-hydroxyacyl-CoA dehydrogenase type-2-Hsd17b10, and 3-ketoacyl-CoA thiolase - Acaa2), where fatty acids are processed to acetyl-CoA, substrate for the TCA cycle [16], [17]. Such a proteome reflects a probable competence of synaptic mitochondria for fatty acid degradation.

**Figure 1.**
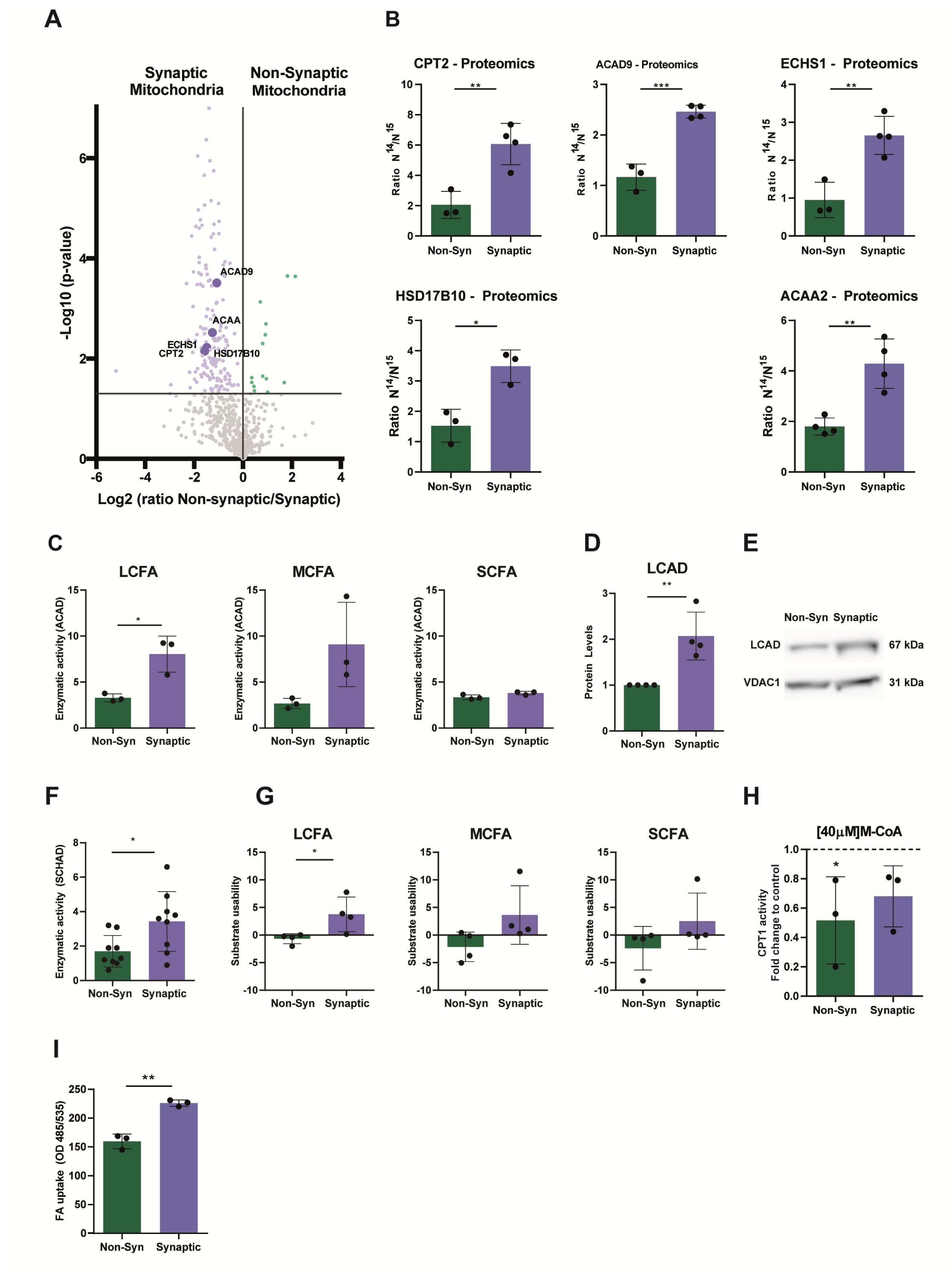
Synaptic mitochondria show higher efficiency towards FAO. (**A**) Volcano plot showing the relative levels of identified proteins in synaptic (Syn) compared to non-synaptic (Non-Syn) mitochondria. The horizontal line defines the p-value statistical significance cutoff (p<0.05). Proteins with differences below the p-value cut-off are depicted in grey. Proteins with increased expression in non-synaptic mitochondria are depicted in red. Proteins with increased expression in synaptic mitochondria are depicted in light blue. Protein involved in FAO are depicted in dark blue. (**B**) Proteomic results of the relative levels comparing synaptic and non-synaptic mitochondria of FAO related proteins highlighted in (A). Data represented as Mean ± SD. n=3-4. Unpaired t-test. *p<0.05; **p<0.01; ***p<0.001. (**C**) Acyl-Coa dehydrogenase (ACAD) activity analysis assessed in isolated synaptic and non-synaptic mitochondria fractions, assessed in the presence of long-(LCFA), medium-(MCFA) and short-chain (SCFA) fatty acids (n=3 per group). Statistics: unpaired Student’s t-test; *p<0,05. (**D**) Quantification via immunoblotting of the expression of LCAD. (**E**) Representative immunoblot images comparing synaptic and non-synaptic mitochondria for LCAD expression depicted in (D). VDAC1 was used as a loading control. Results are presented as fold-change to non-synaptic mitochondria (N=4). Statistics: unpaired Student’s t-test; **p<0,01. (**F**) Short-chain 3-hydroxyacyl-CoA dehydrogenase (SCHAD) activity analysis assessed in isolated synaptic and non-synaptic mitochondria fractions (n=9). Statistics: unpaired Student’s t-test; *p<0,05. (**G**) Metabolism of long-(LCFA), medium-(MCFA) and short-chain (SCFA) fatty acids assessed using Mito Plates S-1^TM^. Results were normalized to wells containing the vehicle control of fatty acids. (N=4). Statistics: unpaired Student’s t-test; *p<0,05. (**H**) CPT1 activity in the presence of 40µM malonyl-CoA (M-CoA) measured in isolated non-synaptic and synaptic mitochondria. Results were normalized to the activity of each fraction without the presence of M-CoA and presented as fold-change. (N=3). Statistics: unpaired Student’s t-test; *p<0,05. (**I**) Fatty acid uptake capacity measured in isolated non-synaptic and synaptic mitochondria. (N=3). Statistics: unpaired Student’s t-test; **p<0,01

### Synaptic mitochondria present higher enzymatic efficiency and higher metabolism of fatty acids when compared to non-synaptic mitochondria

Enzymatic efficiency for FAO via acyl-CoA dehydrogenase (ACAD) activity in synaptic and non-synaptic mitochondria (described in [18], [19]) feed with fatty acids of different chain lengths, revealed a higher ACAD activity in synaptic mitochondria when fueled with long-chain fatty acids (palmitoyl-CoA) (Fig.1C). Complementary immunoblot analysis also showed significantly higher protein levels of acyl-CoA dehydrogenase (LCAD), a protein involved in the processing of long-chain fatty acids in synaptic mitochondria [20] (Fig.1D and E). When furnished with medium-(octanoyl-CoA) and short-chain (butyryl-CoA) fatty acids, no significant differences were observed at the level of ACAD activity (Fig.1C). To further assess the enzymatic efficiency for FAO, we measured the activity of short-chain 3-hydroxyacyl-CoA dehydrogenase (SCHAD, described in [21]), often used to assess fatty acid oxidation capacity as it shows broad activity with 3-hydroxyacyl-CoAs ranging from C4 to C16 [21]. Results obtained reveal a significant increase inSCHAD activity for synaptic mitochondria when compared to their non-synaptic counterparts (Fig.1F). The substrate usability assay using MitoPlates S-1^TM^ measures the ability of mitochondria to metabolize a wide array of substrates, including long-, medium- and short-chain fatty acids. In accordance with the enzymatic assays, we observed significantly higher utilization of long-chain fatty acids by synaptic mitochondria for oxidative metabolism (Fig. 1G). These results indicate that synaptic mitochondria have a more robust enzymatic system for performing fatty acid oxidation (FAO), resulting in a higher metabolic preference for this energy source compared to non-synaptic mitochondria, especially for long-chain fatty acids.

### Synaptic mitochondria are more resistant to FAO inhibition

Mitochondrial fission is essential for axonal colonization and maintenance of synaptic homeostasis [9]. Notably, synaptic mitochondria are also known to be smaller compared to overall neuronal mitochondria [1]. Additionally, different mitochondrial dynamics and morphologies have been associated with different FAO capacities, as mitochondria that undergo fission show reduced malonyl-CoA inhibition of CPT1 [22]. The crosstalk between FAO and mitochondrial dynamics prompted us to examine the activity of CPT1 in the presence of its allosteric inhibitor malonyl-CoA in synaptic and non-synaptic mitochondria. Data obtained showed that non-synaptic mitochondria suffer a significant inhibition of CPT1 activity in the presence of 40µM malonyl-CoA, while synaptic mitochondria showed higher CPT1 activity in the same conditions (Fig.1H). Interestingly, when evaluating the capacity for fatty acid uptake, synaptic mitochondria were also significantly more competent (Fig.1I).

Data obtained in this section shows that synaptic mitochondria have a higher capacity for fatty acid uptake and are simultaneously less sensitive to malonyl-CoA inhibition, which is often associated with smaller, more fissed mitochondria[22].

### FAO increases the respiratory flexibility of brain mitochondria

To interrogate the importance of fatty acid fueling in mitochondrial respiratory flexibility, we performed respiratory measurement assays on synaptic and non-synaptic mitochondria using the Seahorse XFe24 Analyzer, fueling mitochondria with long-(palmitoyl-CoA), medium-(octanoyl-CoA) and short-chain (butyril-CoA) fatty acids (Fig.2A-C) in the presence of relevant mitochondrial substrates pyruvate, succinate and malate. A significant increase in coupling of respiratory complexes (response to ADP, ATP production) and respiratory plasticity (spare capacity, maximum respiration) was observed in both mitochondria populations in the presence of long- and medium-chain fatty acids, with benefits to a higher degree observed in synaptic mitochondria for medium-chain fatty acids (Fig.2D). Concerning short-chain fatty acids, results obtained were not as striking, with synaptic mitochondria showing a significant profit only for ATP production (Fig.2D). Non-synaptic mitochondria showed significant inhibition of basal and maximum respirations as well as in their response to ADP when incubated with short-chain fatty acids (Fig. 2D). These experiments demonstrate that neuronal mitochondria benefit from the respiratory flexibility provided by metabolizing longer-chain fatty acids, even when other standard substrates like pyruvate are available.

**Figure 2.**
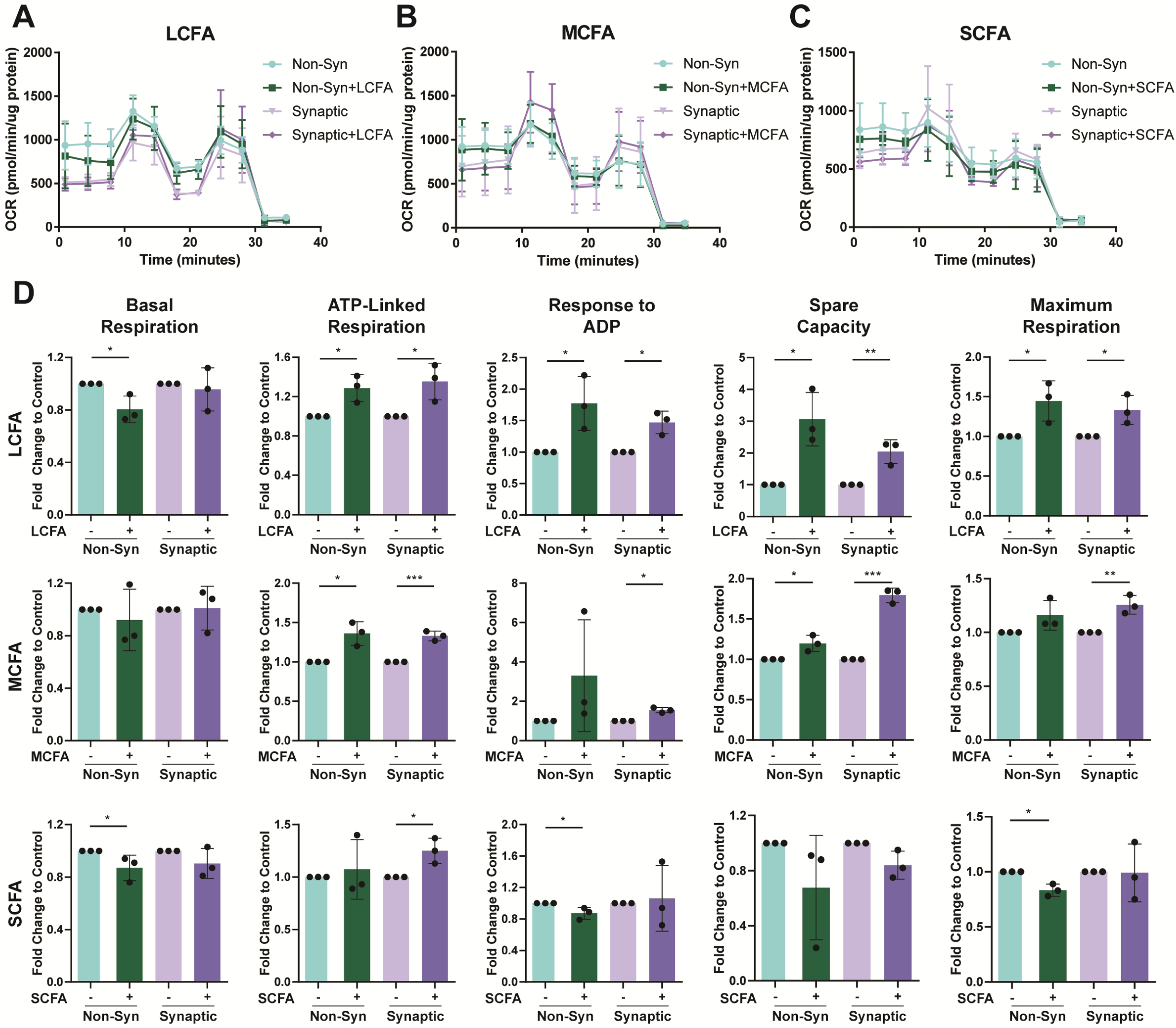
Fatty acid fuelling benefits respiratory flexibility of brain mitochondria. (**A**) Representative graphs of a Seahorse XF Cell Mito Stress Test Kit adapted for isolated mitochondria, performed in a Seahorse XFe24 Analyzer setup in isolated non-synaptic and synaptic isolated mitochondria fractions fuelled with long-(LCFA), medium-(MCFA) and short-chain (SCFA) fatty acids (n=3 per group). (**B**) Respiratory parameters assessed on non-synaptic and synaptic isolated mitochondria fractions in the presence of long-chain fatty acids (LCFA). Results are presented as fold change to the vehicle control of fatty acids (n=3). Statistics: unpaired Student’s t-test; *p<0,05. (**C**) Respiratory parameters assessed on non-synaptic and synaptic isolated mitochondria fractions in the presence of medium-chain fatty acids (MCFA). Results are presented as fold change to the vehicle control of fatty acids (n=3). Statistics: unpaired Student’s t-test; *p<0,05; ***p<0,001. (**D**) Respiratory parameters assessed on non-synaptic and synaptic isolated mitochondria fractions in the presence of short-chain fatty acids (SCFA). Results are presented as fold change to the vehicle control of fatty acids (n=3). Statistics: unpaired Student’s t-test; *p<0,05; **p<0,01.

### Long and medium-chain fatty acid metabolism increases the basal respiration of mouse primary neurons

To explore how fatty acids may affect mitochondrial function in a cellular context, we assessed mitochondrial respiration in whole primary neurons at 10 days in vitro (DIV10) using the Seahorse XFe24 Analyzer. As long- and medium-chain fatty acids were the substrates with most respiratory benefits in brain mitochondria (Fig. 2), we focus our analysis on those two substrates. We observed that in the presence of glucose, treating cells with 25µM of palmitate or octanoate for 4h, as long- and medium-chain fatty acids respectively, significantly increased neuronal basal respiration (Fig.3A,B and C,D, respectively). Interestingly, pre-incubation with the long-chain FAO inhibitor etomoxir before palmitate treatment was enough to return neuronal basal respiration levels to those of the control (Fig.3B). When cells were treated with etomoxir alone, there was a noticeable but not significant reduction in neuronal basal respiration (Fig.3B). The increase in neurons’ basal respiration mediated by octanoate was accompanied by a decrease in spare capacity and maximum respiration (Fig.3D), suggesting the neuronal respiratory fitness better profits from long-chain fatty acid fueling. Importantly, proton leak values associated with possible risk of mitochondrial damage were not increased by either of the fatty acids in question (Supplementary Fig.2A and B). This data shows that both long- and medium-chain fatty acids can benefit neuronal respiration when provided in lower concentrations, particularly for palmitate. Interestingly, inhibiting long-chain FAO was sufficient to deny these effects, showing that long-chain fatty acids can boost the flexibility of neuronal respiration even in the presence of glucose.

**Figure 3.**
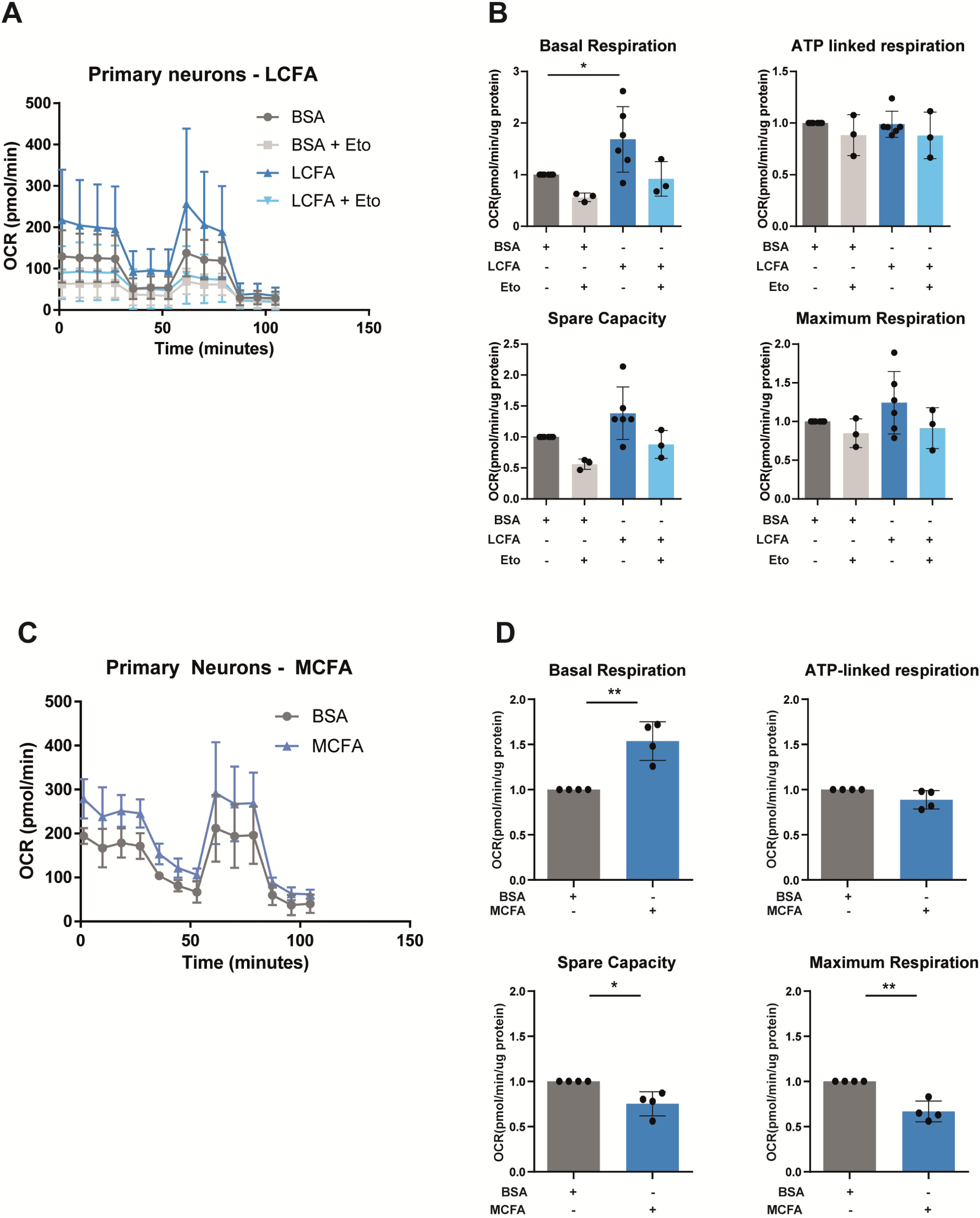
Neuronal respiration benefits from FAO. (**A**) Representative graphs of a Seahorse XF Cell Mito Stress Test Kit, performed in a Seahorse XFe24 Analyzer setup in primary neurons (DIV10) fueled with long-chain fatty acids (LCFA) in the form of Palmitate (n=3-6 per group). (**B**) Analysis of respiratory parameters of primary neurons fueled with Palmitate. Results are presented as fold change to the vehicle control of fatty acids (BSA). (n=3-6 per group). Statistics: ordinary one-way ANOVA with Dunnet’s multiple comparisons test; *p<0,05. (**C**) Representative graphs of a Seahorse XF Cell Mito Stress Test Kit, performed in a Seahorse XFe24 Analyzer setup in primary neurons (DIV10) fueled with medium-chain fatty acids (MCFA) in the form of Octanoate (n=4 per group). (**D**) Analysis of respiratory parameters of primary neurons fueled with Octanoate. Results are presented as fold change to the vehicle control of fatty acids (BSA). (n=4 per group). Statistics: unpaired Student’s t-test; *p<0,05; **p<0,01.

### Fatty acid fueling significantly contributes to neuronal ATP levels

Following the results from the respiratory assays described in the previous section, we aimed to understand the effect of long- and medium-chain fatty acid oxidation on neuronal ATP levels. Mouse primary neurons (DIV10) were treated with palmitate and octanoate for 4h in the presence of glucose and ATP levels were quantified. Fueling primary neurons with palmitate did not affect overall neuronal ATP levels, however when cells were pre-incubated with etomoxir, a significant reduction was observed (Fig.4A). For octanoate oxidation, we also observed a significant increase in primary neuron ATP levels upon medium-chain fatty acid fueling (Fig.4B). Data obtained shows that even in the presence of glucose, preventing long-chain FAO is detrimental to the maintenance of ATP levels in neurons, suggesting a role for FAO in the energetic homeostasis of these cells. At the same time, medium-chain FAO can significantly contribute to the increase in the pool of energy available to neurons.

**Figure 4.**
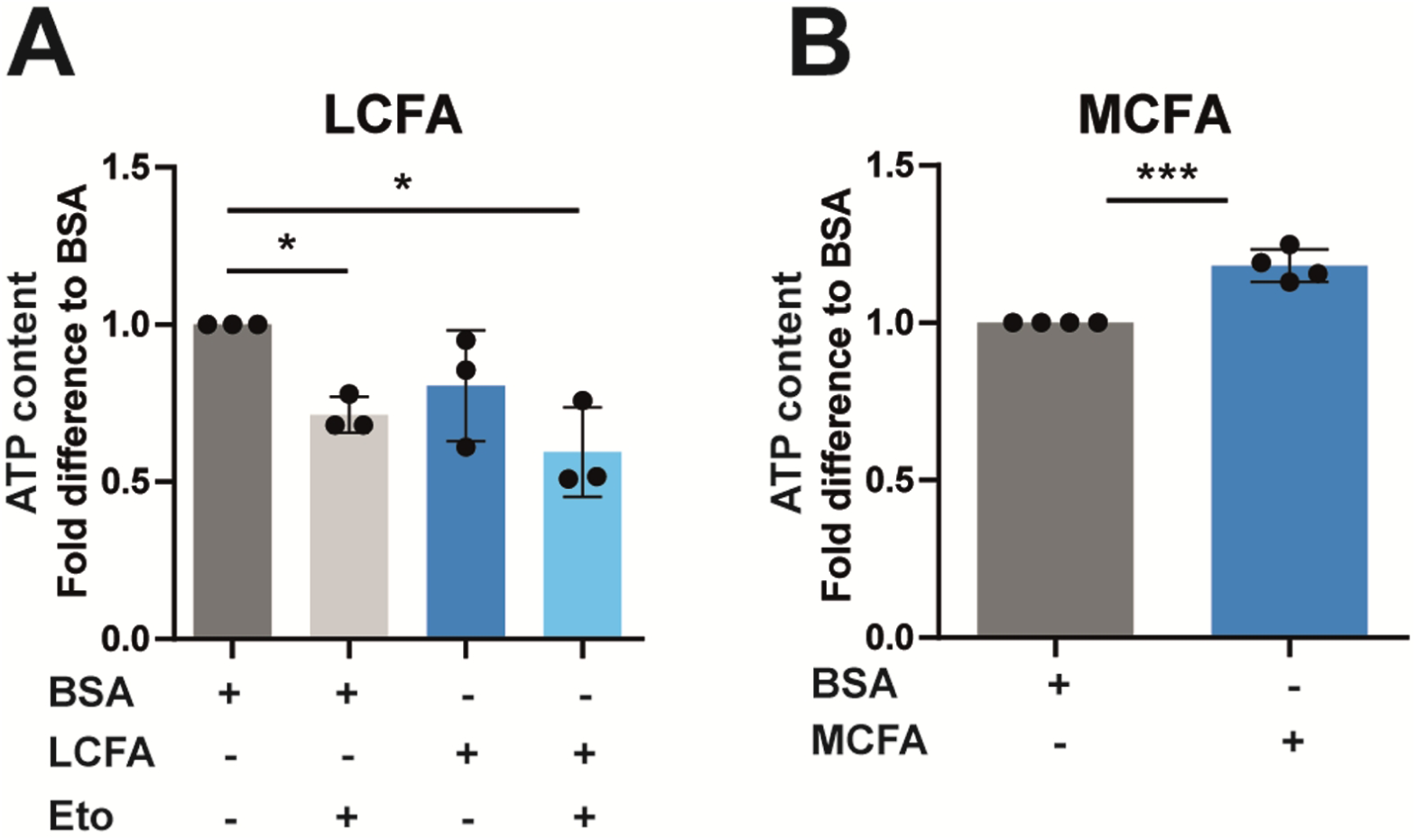
Fatty acid fueling significantly contributes to neuronal ATP levels. (**A**) Analysis of the effect of Palmitate and Etomoxir and (**B**) Octanoate on the ATP content of primary neurons (DIV10). Results are presented as fold change to the vehicle control of fatty acids (BSA). (n=4 per group). Statistics: (Palmitate and Etomoxir) ordinary one-way ANOVA with Dunnet’s multiple comparisons test; *p<0,05; (Octanoate) unpaired Student’s t-test; ***p<0,001 for Octanoate.

### Neurons naturally uptake fatty acids at the periphery in the presence of glucose

As the role of glucose is well known to maintain neuronal homeostasis, evidence starts to arise showing that in the absence of this fuel, fatty acids derived from lipid droplets can sustain electrical activity at neuronal periphery [23]. Nevertheless, to our knowledge, nobody has yet shown if fatty acid uptake occurs specifically at the synapse in the presence of glucose. To answer this, we cultured mouse primary neurons in microfluidic devices [24], allowing isolation of axons from the somal region [15]. In this setup, and after 10 days *in vitro* (DIV10), a 24h pulse of BODIPY 558/568 (Red-C12) was delivered the axonal compartment, and its uptake was assessed at both neuronal (Tubulin β3) and mitochondrial levels (HSP60). Indeed, we observed the uptake of Red-C12 labeled fatty acids uptake at neuronal periphery (Fig.5A). Moreover, some Red-C12 fatty acids were also found near axonal mitochondria, suggesting that even in the presence of glucose fatty acids serve as fuels for axonal mitochondria (Fig.5B). Importantly, exclusive peripheric uptake of fatty acids was confirmed by the absence of signal in the somal compartment (Supplementary Fig.3A). Fuel uptake at a whole cellular level was also observed when Red-C12 was delivered to neurons plated in coverslips (Supplementary Fig.3B).

**Figure 5.**
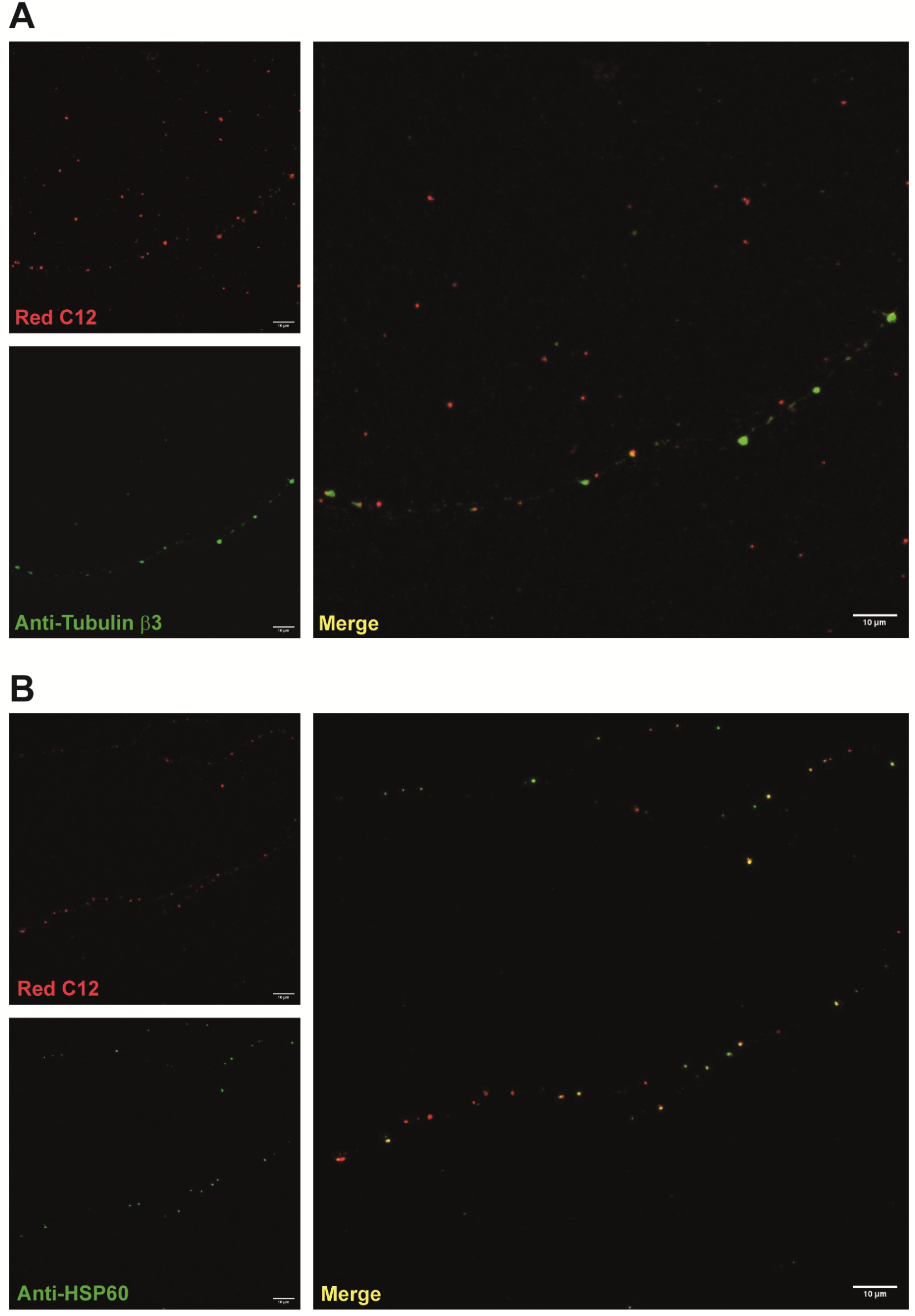
Fatty acid uptake occurs at neuronal periphery in the presence of glucose. Confocal images show uptake of Red-C12 incubated for 24h exclusively in the axonal compartment of mouse primary neurons (DIV10) cultured in microfluidic devices. Fatty acid uptake was observed at the (**A**) cellular (via co-staining with neuronal marker anti-Tubulin β3 antibody) and (**B**) mitochondrial (via co-staining with mitochondrial marker anti-HSP60 antibody) levels, evidence showing that metabolization of this fuel can occur locally at the periphery of neurons.

### Primary neurons show increased pre-synaptic activity upon palmitate fueling

Mitochondrial ATP supply at the synapse is well documented and is essential for the maintenance of proper firing activity [25], [26]. Although the contribution of glucose has been well documented in this extent [25], [27], the question remains if neuronal activity may also benefit from additional fuels such as fatty acids. To assess this, we performed whole-cell patch-clamp recordings in primary neurons (DIV 19-21) treated with 25µM of palmitate or its vehicle control BSA for 4h in complete medium containing glucose (Fig.6A). Spontaneous postsynaptic currents were evaluated, since changes in amplitude reflect postsynaptic alterations, while changes in frequency indicate modifications in presynaptic function or synapse number. Results obtained revealed a significant impact of palmitate in neuronal activity, showing increased current frequency when compared with neurons treated with BSA (Fig.6B). Moreover, incubating primary neurons with etomoxir prior to adding palmitate was sufficient to reduce the frequency of spontaneous postsynaptic currents to control values, thereby confirming the specificity of the effect of long-chain fatty acid oxidation in synaptic activity (Fig.6B). No differences in current amplitude were observed in any of the treatments performed (Fig.6C), as well as in the expression of glutamate receptors, such as NMDAR2A/B and GluA1 (Fig.6D and E). This data suggests that neurons have the flexibility to utilize different fuel sources even in the presence of glucose. The ability to oxidize energy-rich fuels like palmitate may be particularly important in synapses, where there is a constant high demand for energy.

**Figure 6.**
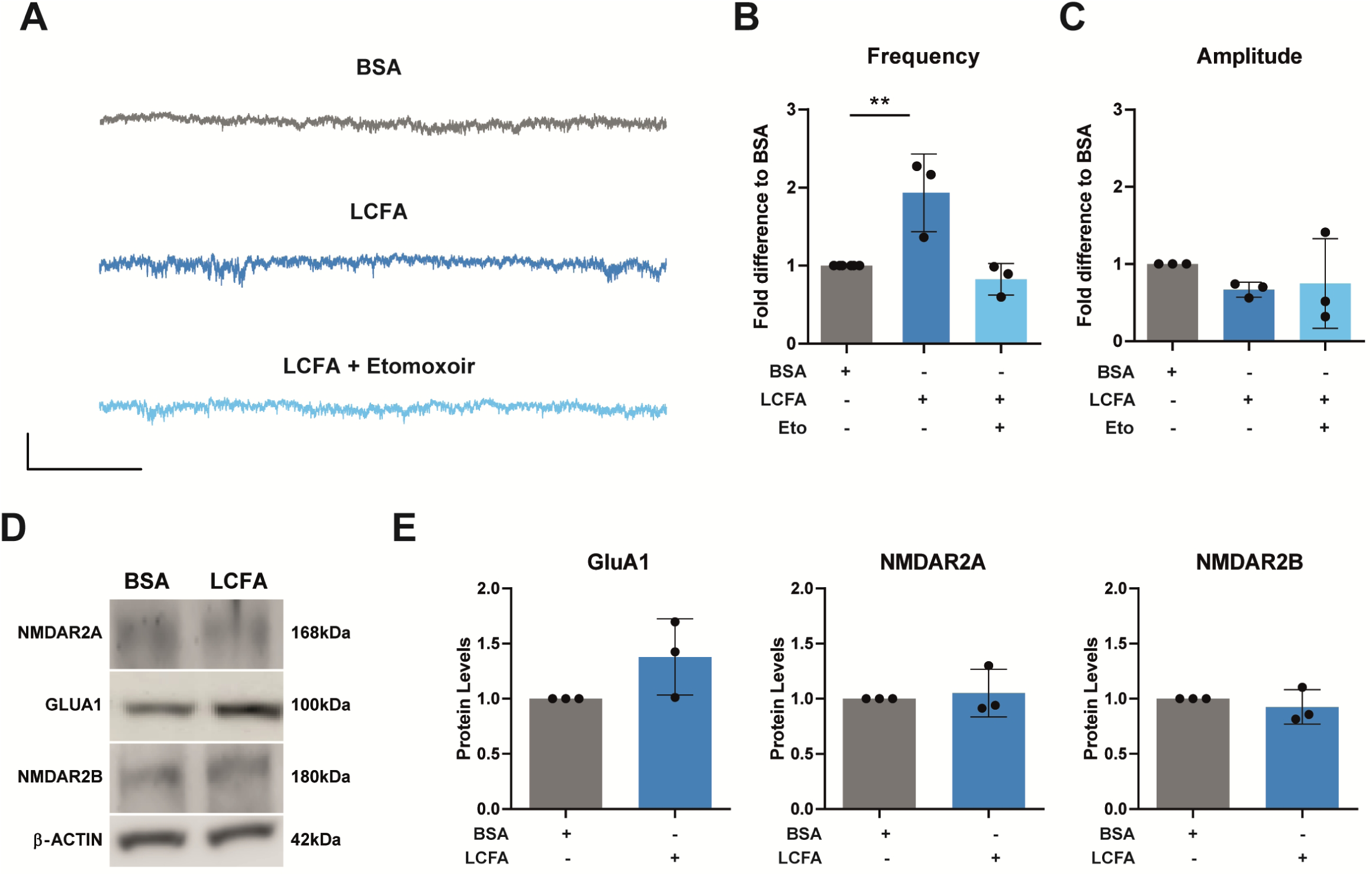
Long-chain FAO increases synaptic potentiation. (**A**) Representative traces of spontaneous postsynaptic currents in primary neurons (DIV19-21) incubated with long-chain fatty acids (LCFA), alone or in combination with the inhibitor Etomoxir, as well as vehicle control BSA. Graphical analysis of whole-cell patch-clamp recording depicting frequency **(B)** and amplitude (**C**) of spontaneous postsynaptic currents performed in primary neurons (DIV19-21) fueled with long-chain fatty acids (LCFA) in the form of Palmitate. Etomoxir was incubated prior to palmitate to assess specificity of palmitate Effect. Recording conducted in Axopatch 200B (Axon Instruments) amplifier. Results are presented as fold-change to BSA (N=3). Statistics: ordinary one-way ANOVA with Dunnet’s multiple comparisons test; **p<0,01 (**D**) Representative immunoblot images comparing primary neurons (DIV19-21) fueled with palmitate or BSA for the expression of GluA1, MDAR2A and NMDAR2B. B-actin was used as a loading control. Statistics: unpaired Student’s t-test. (E) Quantification via immunoblotting of the expression of GluA1, NMDAR2A and NMDAR2B. Results are presented as fold-change to BSA (N=3).

## DISCUSSION

The long-held dogma that the brain does not rely on FAO is undergoing increasing scrutiny, with research showing that this fuel source can contribute to total brain oxidative energy production [28], astrocytic metabolism [29] and metabolic coupling between neurons and glia [14], [30]. Recent evidence shows that, in the absence of glucose, fatty acids can be stored in lipid droplets in axons, and its flux to mitochondria for constant degradation is mediated by the DDHD2 lipase, which is why historically it has been assumed that the presence of lipid droplets in neurons is neglectable [23]. Indeed, DDHD2 loss of function caused lipid droplet accumulation in axons, leading to significant cognitive impairment in humans and mice, advocating the importance of functional metabolism of fatty acids to preserve neuronal health [23], [31], [32]. Nevertheless, evidence for the usage of fatty acids in neuronal and synaptic homeostasis under conditions where glucose is still present remains to be assessed.

In this extent, our results provide solid proof that in the presence of glucose, FAO significantly contributes to neuronal homeostasis. We have shown for the first time that fueling neurons with concentrations of palmitate that do not cause lipotoxicity significantly benefits excitatory synaptic activity even in the presence of glucose, which was rescued to basal levels by inhibiting the carnitine shuttle import of palmitate with etomoxir. We observe the same outcome in oxygen consumption assays, whereby fueling neurons with palmitate significantly benefited cellular respiration, while pre-incubation with etomoxir reversed this phenotype. This demonstrates that long-chain fatty acid oxidation (LCFAO) can fuel synaptic activity even in the presence of glucose, challenging the view that glucose is the sole metabolic substrate contributing to maintain homeostasis at neuronal periphery. The fact that we were able to observe the uptake of Red-C12 fatty acids at neuronal periphery in the presence of glucose, and its location in the vicinity of axonal mitochondria, also adds to the view that FAO is a physiologically relevant process at the synapse. Interestingly, neurons incubated with etomoxir experienced a significant decrease in ATP levels, showing their need to resort to FAO even in the presence of glucose to maintain their energy balance. This goes in line with the research from Panov and colleagues that showed that combining fatty acids with either pyruvate, glutamate or succinate increased rates of brain mitochondrial respiration in all metabolic states [12].

The increase in pre-synaptic activity and overall respiratory benefits upon palmitate fueling observed may be linked with an increased specialization of synaptic mitochondria in dealing with this fuel. Our data showed synaptic mitochondria’s flexibility towards FAO is reflected in higher competence in oxidizing fatty acid of longer chain-lengths, with increased activity of β-oxidation enzymes, higher metabolism and capacity for uptake, and less proneness for inhibition via Malonyl-CoA when compared with non-synaptic mitochondria. Moreover, long- to medium-chain fatty acid fueling significantly increased the flexibility of mitochondrial electron transport chain even in the presence of other fuels such as pyruvate. To our knowledge, no studies have evaluated the fuel preference of synaptic mitochondria; however, specific features such as their smaller size and fission dynamics may indicate higher capacity for FAO. A study in HepG2 cells and primary rat hepatocytes from Ngo and colleagues showed a strong correlation between mitochondrial fragmentation and increased FAO rates, as inducing fragmentation led to reduced malonyl-CoA inhibition of CPT1, specifically increasing long-chain FAO [22]. Our data corroborated this view, as we showed that synaptic mitochondria are more resistant to malonyl-CoA inhibition than non-synaptic mitochondria, which implies that the flow of importing long-chain fatty acids is less prone to disruption. The preference of synaptic mitochondria for degrading fatty acids of higher chain-lengths was strengthened by our enzymatic and substrate usability data, revealing increased enzymatic activity and metabolism in the presence of this fuel. Although the importance of glucose in neuronal and synaptic function is well established [33], the energetic demands and pressures of the environment surrounding synaptic terminals raises the question of whether additional fuel sources might be beneficial for maintaining proper synaptic circuitry function. Astrocytes, for instance, have shown remarkable flexibility in substrate utilization, demonstrating competence in both glycolysis and fatty acid oxidation (FAO) [14], [34]. While they can remove peroxidized lipids that could potentially damage neurons, they also provide lactate to fuel synaptic plasticity [35], [36]. Advances in the understanding of astrocyte metabolism have shown also that medium-chain fatty acid (MCFA) fueling promotes astrocytic ketogenesis [37], suggesting that astrocytes could potentially supply additional fuels, such as ketone bodies or even fatty acids, to the synapse. Altogether, our work shows that the fueling preferences of synaptic mitochondria allow them to deal with energy-rich substrates like fatty acids, which in combination with glucose serve as hubs to sustain neuronal and synaptic homeostasis. Considering that synaptic dysfunction typically precedes neuronal loss, and that metabolic dysfunctions are a key factor in the etiology of brain diseases such as Alzheimer’s (AD), Parkinson’s (PD), and amyotrophic lateral sclerosis (ALS) [38], [39], dissecting the metabolic fingerprint of synaptic terminals could be crucial for understanding which pathways become disrupted in neurodegeneration.

## CONCLUSION

Our work unravels for the first time the fuel preference and flexibility of synaptic mitochondria. Synaptic mitochondria play a crucial role in supplying neurons with ATP to maintain proper protein synthesis and synaptic vesicle cycling, as well as a healthy neuronal circuitry [25], [40]. Further unravelling synaptic fuel preferences, particularly in a firing state, will be key to understand the early stages of neurodegenerative diseases and ultimately prevent its progression.

## ACKNOWLEDGEMENTS

We would like to thank Bart de Strooper and Kathleen Craessaerts (VIB, Leuven, Belgium) for their initial support to this project and for sharing fruitful scientific discussions. We acknowledge also Jeffrey Savas and his team for all the help with performing and analysing the proteomics data. We thank the members of the VMorais Lab for their support and feedback. We would also like to thank the BioImaging Facility and the Rodent Facility of Gulbenkian Institute for Molecular Medicine for their technical support, and we also acknowledge the funding PPBI-POCI-01-3700145-FEDER-022122.

## AUTHOR CONTRIBUTIONS

Conceptualization: B.C.A. and V.A.M.; methodology: B.C.A., A.F.P., J.G.R., S.C.P., S.H.V.; formal analysis: B.C.A. and V.A.M.; investigation: B.C.A. and V.A.M.; writing – original Draft: B.C.A.; writing – review and editing: B.C.A. A.F.P. and V.A.M.; funding acquisition, V.A.M.; supervision: V.A.M.

All authors have read and agreed to the published version of the article.

## SUPPLEMENTARY FIGURE TITLES AND LEGENDS

**Supplementary Figure 1.**
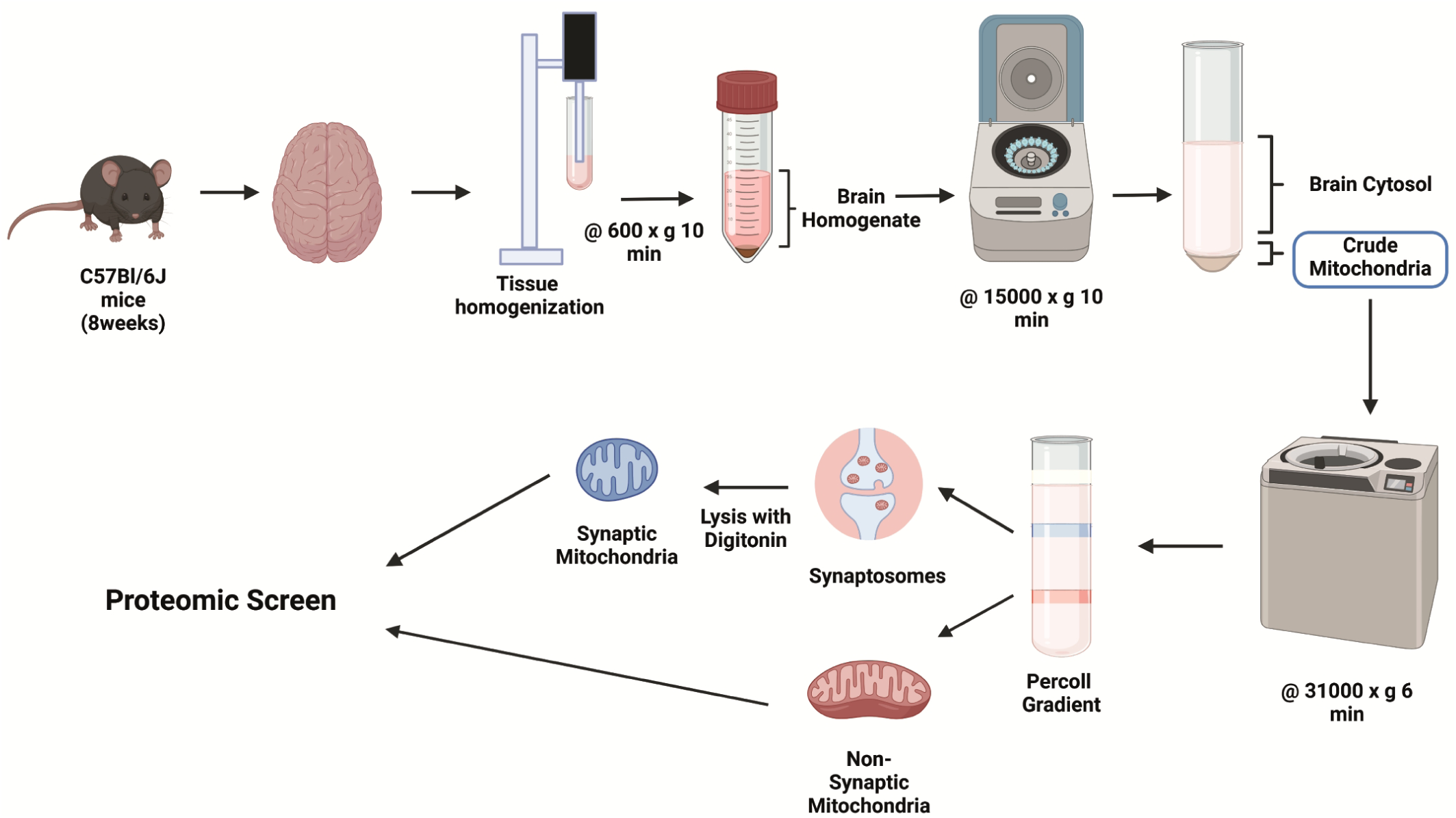
The proteome of synaptic mitochondria shows higher affinity for FAO. Representative scheme of the isolation protocol of synaptic and non-synaptic mitochondria from the brains of 8 week-old C57Bl/6J mice.

**Supplementary Figure 2.**
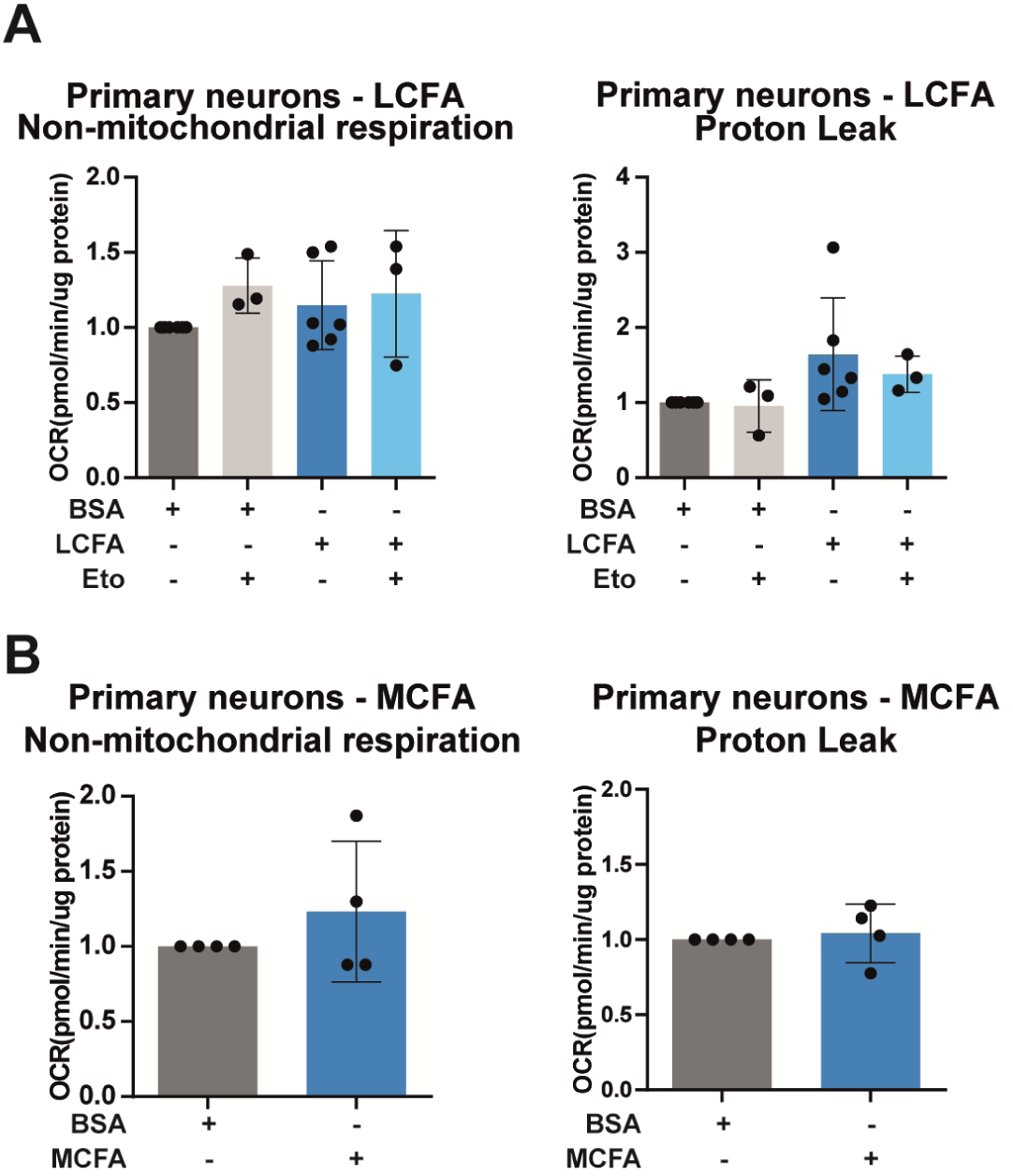
Integrity of the ETC of neuronal mitochondria is maintained upon fatty acid fueling. (**A**) Analysis of respiratory parameters of primary neurons (DIV10) with long-chain fatty acids (LCFA) in the form of Palmitate under a Seahorse XF Cell Mito Stress Test Kit, performed in a Seahorse XFe24 Analyzer setup. Results are presented as fold change to the vehicle control of fatty acids (BSA). (n=3-6 per group). Statistics: ordinary one-way ANOVA with Dunnet’s multiple comparisons test. (**B**) Analysis of respiratory parameters of primary neurons (DIV10) with medium-chain fatty acids (MCFA) in the form of Octanoate under a Seahorse XF Cell Mito Stress Test Kit, performed in a Seahorse XFe24 Analyzer setup. Results are presented as fold change to the vehicle control of fatty acids (BSA). (n=4 per group; Statistics: unpaired Student’s t-test.

**Supplementary Figure 3.**
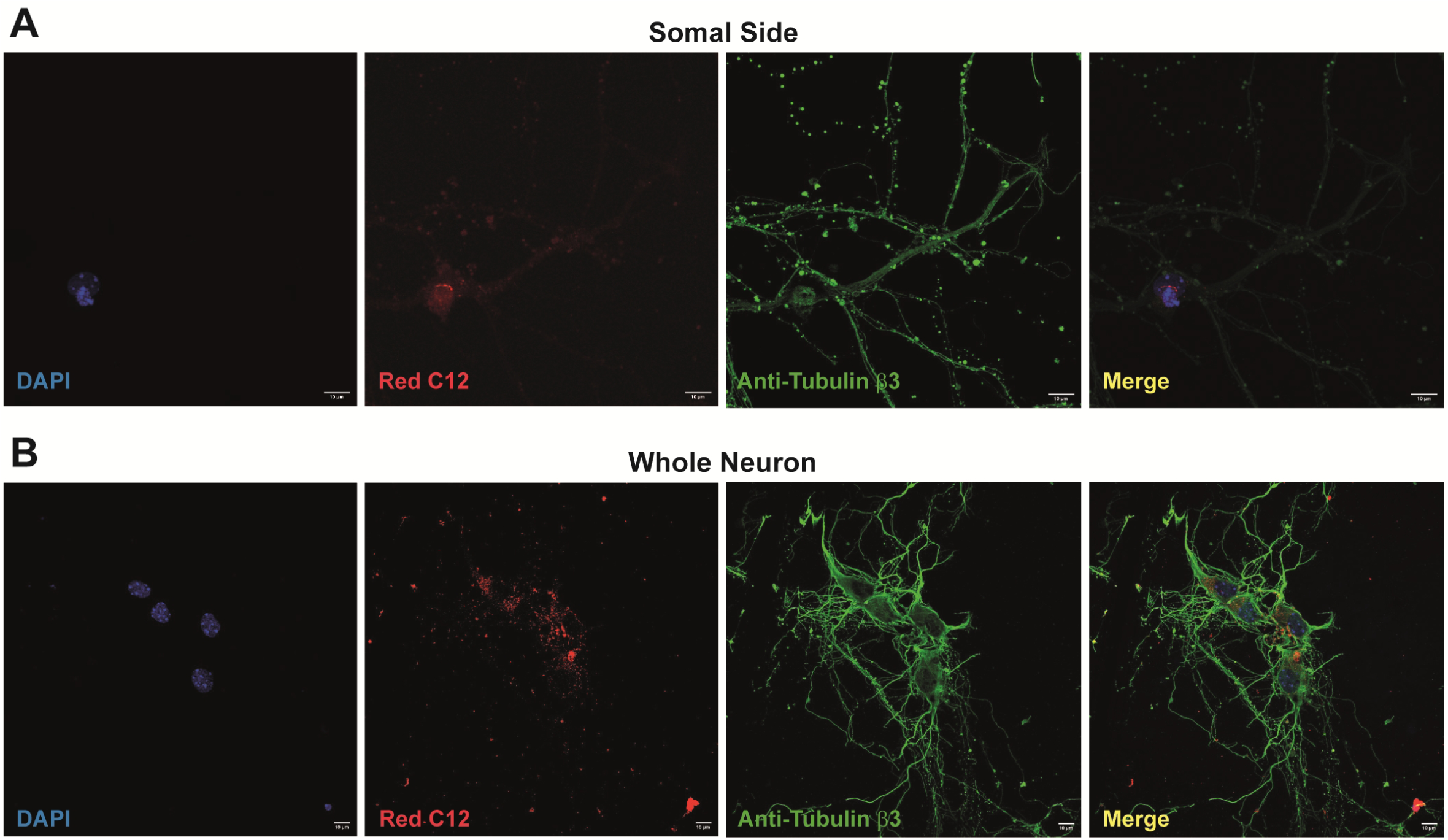
Neurons are able to uptake fatty acids in the presence of glucose. (**A**) Confocal images show absence of signal of Red-C12 in the somal compartment of microfluidic devices after a 24h incubation period exclusively in the axonal compartment. (**B**) Confocal images show uptake of Red-C12 incubated 24h in mouse primary neurons (DIV10). Fatty acid uptake was observed at the cellular level via co-staining with neuronal marker anti-Tubulin β3 antibody.

## STAR★Methods

### Key resources table

| REAGENT or RESOURCE | SOURCE | IDENTIFIER |
| --- | --- | --- |
| **Antibodies** |  |  |
| Anti-LCAD | Proteintech | 17526-1-AP; AB_2219661 |
| Anti-VDAC1 | Abcam | ab154856; RRID: AB_2687466 |
| Anti-NMDAR2A | Abcam | ab169873 |
| Anti-NMDAR2B | Abcam | ab65783 |
| Anti-GLUA1 | Millipore | AB1504; AB_2113602 |
| Anti-β-Actin | Sigma-Aldrich | A5441 |
| Anti-HSP60 | BD Biosciences | 611563; RRID: AB_399009 |
| Anti-β3 tubulin | BioLegend | 802001; RRID: AB_2564645 |
| Alexa Fluor 488 Anti-Rabbit | Invitrogen | A21206; RRID: AB_2535792 |
| Alexa Fluor 488 Anti-Mouse | Invitrogen | A21202; AB_141607 |
| HRP Conjugated Anti-Mouse | Bio-Rad | 1706516; RRID: AB_2921252 |
| HRP Conjugated Anti-Rabbit | Bio-Rad | 1706515; RRID: AB_11125142 |
| **Chemicals, peptides, and recombinant proteins** |  |  |
| DAPI | Sigma-Aldrich | D9542 |
| Paraformaldehyde (PFA) | Electron Microscopy Sciences | 15710 |
| Bovine Serum Albumin (BSA) | Sigma-Aldrich | A9647 |
| Hank's Balanced Salt Solution (HBSS) | Gibco | 14175095 |
| HEPES | Gibco | 15630056 |
| Trypsin | Gibco | 15090046 |
| DNase I | Sigma-Aldrich | DN25 |
| Horse Serum (HS) | Gibco | 2605088 |
| Minimum Essential Medium (MEM) | Gibco | 31095029 |
| Glucose | Gibco | A2494001 |
| Penicillin-Streptomycin | Gibco | 15140122 |
| NeuroBasal medium (NB) | Gibco | 21103049 |
| Poly D-Lysine | Sigma-Aldrich | P7280 |
| L-Glutamine | ThermoScientific | 25030024 |
| B-27 | Gibco | 17504044 |
| VectaShieldAntifade Mounting Medium | Vector laboratories | H-1000 |
| Sucrose | Sigma-Aldrich | 84100 |
| D-Mannitol | Sigma-Aldrich | M4125 |
| EGTA | Sigma-Aldrich | E4378 |
| Percoll | Sigma-Aldrich | GE17-0891-02 |
| Digitonin | Sigma-Aldrich | D141 |
| Protease Inhibitor | ThermoScientific | 78443 |
| Malate | Sigma-Aldrich | M1000 |
| Succinate | Sigma-Aldrich | S2378 |
| Sodium Pyruvate | ThermoScientific | LTID 11360-039 |
| Potassium Phosphate Dihydrogen (KH<sub>2</sub>PO<sub>4</sub>) | Sigma-Aldrich | P5379 |
| Di-Potassium Phosphate (K<sub>2</sub>HPO<sub>4</sub>) | Sigma-Aldrich | 795496 |
| Potassium Cyanide (KCN) | VWR | 26802.234 |
| Etomoxir | Sigma-Aldrich | E1905 |
| Rotenone | Sigma-Aldrich | R8875 |
| Antimycin A | Sigma-Aldrich | A8674 |
| FCCP | Sigma-Aldrich | C2920 |
| Cytochrome C | Sigma-Aldrich | C7752 |
| Phenazine methosulfate | Sigma-Aldrich | P9625 |
| MOPS | Sigma-Aldrich | M1254 |
| DTT (Dithiotreitol) | Biochemica | A1101,0025 |
| Triton X100 | Sigma-Aldrich | T8787 |
| NADH | Sigma-Aldrich | N8129 |
| N-ethylmaleimide | Sigma-Aldrich | E3876 |
| Acetoacetyl coenzyme A sodium salt hydrate | Sigma-Aldrich | A1625 |
| Palmitoyl coenzyme A lithium salt | Sigma-Aldrich | P9716 |
| Octanoyl coenzyme A lithium salt | Sigma-Aldrich | O6877 |
| Butyryl coenzyme A lithium salt hydrate | Sigma-Aldrich | B1508 |
| DTNB (5 5'-Dithiobis(2-Nitrobenzoic Acid) | Sigma-Aldrich | D8130 |
| EDTA | Sigma-Aldrich | E5134 |
| Malonyl coenzyme A | Sigma-Aldrich | M4263 |
| Carnitine Hydrochloride | Sigma-Aldrich | C0283 |
| BODIPY™ 558/568 C12 | Invitrogen | D3835 |
| **Critical commercial assays** |  |  |
| Fatty Acid Uptake Probe | Dojindo | UP07 |
| ECL Western Blotting reagents | Cytiva | RPN2106 |
| Pierce™ BCA protein assay kit | ThermoScientific | 23225 |
| Bio-Rad protein assay | Bio-Rad | 5000001 |
| Seahorse XF Palmitate-BSA FAO Substrate | Agilent Technologies | HPA102720100 |
| Luminescent ATP Detection Assay Kit | Abcam | ab113849 |
| **Deposited data** |  |  |
| Proteomics Data | MassIVE | MSV000092568 |
| **Experimental models: Organisms/strains** |  |  |
| Mouse: C57BL/6J | Mice bred in-house | N/A |
| **Software and algorithms** |  |  |
| Prism v8.4.3 | Graphpad | https://www.graphpad.com/ |
| Fiji (Image J) |  | https://imagej.net/software/fiji/ |
| Adobe Illustrator |  | https://www.adobe.com/ |
| **Other** |  |  |
| MitoPlates S-1 | Biolog | 14105 |
| Microfluidics 450 mm microgroove barrier | Xona | SND450 |
| Zeiss LSM 880 Confocal microscope | Zeiss | N/A |
| Amersham 680 RGB | GE Healthcare | N/A |
| Amersham ImageQuant 800 | GE Healthcare | N/A |
| Mini Gel Tank and Blot Module Set | Invitrogen | NW2000 |
| Potter Elvehjem 30mL | Fisher Scientific | 10669922 |
| BioTek Synergy HTX Multimode Reader | BioSPX |  |
| Seahorse XF24e Analyzer | Agilent Technologies | S7801B |

### Experimental model and study participant details

#### Animals

C57BL/6J mice were obtained and housed in the Rodent Facility from Gulbenkian Institute for Molecular Medicine (GIMM), in a temperature-controlled room at 20-24°C. Stud males were housed individually. Females were housed in groups of 2-5 per cage and synchronization of their estrous cycles was achieved by exposing them to male’s urine 48h prior to mating. Whole brains from embryos at embryonic day 18 (E18) were used for the establishment of primary neuron cultures. All the procedures were approved by the Portuguese National Authority for Animal Health (DGAV), as well as by the institute’s animals’ well-being office (ORBEA-iMM). The study was carried out in compliance with the ARRIVE guidelines [15].

### Methods details

#### Isolation of synaptic and non-synaptic mitochondria from mouse brain

Brains were isolated from 8 weeks old C57BL/6J female mice and synaptic and non-synaptic mitochondrial enriched fraction was performed as previously described [16]. Briefly, tissue homogenization was performed in cold isolation buffer (IB) containing 10mM HEPES, 225mM sucrose, 75mM D-Mannitol and 1mM EGTA, pH 7.4 followed by a centrifugation at 600xg, 10min, 4°C. Supernatant was then collected and further centrifuged at 15000xg, 10min, 4°C, resuspended in 15% Percoll and applied on an ascending Percoll gradient of 40% and 24%. Synaptosomes (in the interphase 15% - 24% Percoll) and non-synaptic mitochondria (in the interphase 24% - 40% Percoll) were collected and resuspended in cold IB and centrifuged at 20,000xg and 10,000xg for 10min, 4°C. To collect synaptic mitochondria, synaptosomes were disrupted with 0,6mg/mL of digitonin for 15min in agitation, 4°C, followed by two centrifugations at 10,000xg, 10min, 4°C. The protein content of isolated brain mitochondria fraction samples was measured using Pierce™ BCA Protein Assay Kit. Isolated mitochondrial fractions were used fresh or stored at -80° until further use.

#### Proteomics

The stable 15N isotope labelling of mammals was accomplished essentially as previously described.52,53 Briefly, FVB mice of both sexes were purchased immediately after weaning. After a week-long acclimation, mice were placed on ad libitum 15Ndiet (MF-SpirulinaN-IR; Cambridge Isotopes). For the dynamic labelling experiments (single-generation), mice stayed on the 15N diet for 4 months and were sacrificed. Mice were anesthetized with 3% isoflurane followed by acute decapitation. Brain tissues were harvested, flash frozen in a dry ice/ethanol bath, and stored at -80°C. For consequent MS studies, brain homogenates from naturally occurring N14 fresh brains from C57BL/6J female mice were mixed in a 1:1 ratio with brain homogenates coming from N15 labelled frozen brains. From the N14/N15 mixed brain homogenate, mitochondria were isolated as described above.

#### MS sample preparation

Mitochondrial samples were homogenized in the lysis buffer (0.1 M Tris-HCl, pH 7.6, 20% SDS and 1 M DTT with 13 protease inhibitor cocktail. 54 The mixtures were incubated at 95°C for 5 min. The samples were then sonicated for 10 min and centrifuged at 16,100 x g for 10 min. The supernatant was collected and the proteins were precipitated by the methanol/chloroform method. Protein pellets were resuspended in 8M urea prepared in 100 mM ammonium bicarbonate solution and processed with ProteaseMAX according to the manufacturer’s protocol. The samples were reduced with 5 mM Tris(2-carboxyethyl)phosphine (TCEP; vortexed for 1 hour at RT), alkylated in the dark with 10 mM iodoacetamide (IAA; 20 min at RT), diluted with 100 mM ABC, and quenched with 25 mM TCEP. Samples were diluted with 100 mM ammonium bicarbonate solution, and digested with Trypsin (1:50,) for overnight incubation at 37 C with intensive agitation. The next day, reaction was quenched by adding 1% trifluoroacetic acid (TFA). The samples were desalted using Peptide Desalting Spin Columns. All samples were vacuum centrifuged to dry.

#### Tandem mass spectrometry

Three micrograms of each sample were auto-sampler loaded with a Thermo EASY nLC 100 UPLC pump onto a vented Acclaim Pepmap 100, 75 um x 2 cm, nanoViper trap column coupled to a nanoViper analytical column (3 mm, 100 A°, C18, 0.075 mm, 500 mm) with stainless steel emitter tip assembled on the Nanospray Flex Ion Source with a spray voltage of 2000 V. An Orbitrap Fusion was used to acquire all the MS spectral data. Buffer A contained 94.785% H2O with 5% ACN and 0.125% FA, and buffer B contained 99.875% ACN with 0.125% FA. The chromatographic run was for 4 hours in total with the following profile: 0-7% for 7, 10% for 6, 25% for 160, 33% for 40, 50% for 7, 95% for 5 and again 95% for 15 mins receptively. Samples were injected and analyzed by three or six times. We used CID-MS2 method for these experiments as previously described.55 Briefly, ion transfer tube temp = 300°C, Easy-IC internal mass calibration, default charge state = 2 and cycle time = 3 s. Detector type set to Orbitrap, with 60K resolution, with wide quad isolation, mass range = normal, scan range = 300-1500 m/z, max injection time = 50 ms, AGC target = 200,000, microscans = 1, S-lens RF level = 60, without source fragmentation, and datatype = positive and centroid. MIPS was set as on, included charge states = 2-6 (reject unassigned). Dynamic exclusion enabled with n = 1 for 30 s and 45 s exclusion duration at 10 ppm for high and low. Precursor selection decision = most intense, top 20, isolation window = 1.6, scan range = auto normal, first mass = 110, collision energy 30%, CID, Detector type = ion trap, OT resolution = 30K, IT scan rate = rapid, max injection time = 75 ms, AGC target = 10,000, Q = 0.25, inject ions for all available parallelizable time.

#### MS data analysis and quantification

Protein identification/quantification and analysis were performed with Integrated Proteomics Pipeline - IP2 (Bruker, Madison, WI. http://www.integratedproteomics.com/) using ProLuCID,56,57 DTASelect2,58,59 Census and Quantitative Analysis. Spectrum raw files were extracted into MS1, MS2 files using RawConverter (http://fields.scripps.edu/downloads.php). The tandem mass spectra (raw files from the same sample were searched together) were searched against UniProt mouse (downloaded on 03-25-2014) protein databases60 and matched to sequences using the ProLuCID/SEQUEST algorithm (ProLuCID version 3.1) with 50 ppm peptide mass tolerance for precursor ions and 600 ppm for fragment ions. The search space included all fully and half-tryptic peptide candidates within the mass tolerance window with no-miscleavage constraint, assembled, and filtered with DTASelect2 through IP2. To estimate protein probabilities and false-discovery rates (FDR) accurately, we used a target/decoy database containing the reversed sequences of all the proteins appended to the target database.60 Each protein identified was required to have a minimum of one peptide of minimal length of six amino acid residues; however, this peptide had to be an excellent match with an FDR < 1% and at least one excellent peptide match. After the peptide/spectrum matches were filtered, we estimated that the protein FDRs were%1% for each sample analysis. Resulting protein lists include subset proteins to allow for consideration of all possible protein forms implicated by at least two given peptides identified from the complex protein mixtures. Then, we used Census and Quantitative Analysis in IP2 for protein quantification. Static modification: 57.02146 C for carbamidomethylation. Quantification was performed by the built-in module in IP2.

#### Acyl-CoA dehydrogenase assay

The activity of Acyl-CoA dehydrogenase was measured in non-synaptic and synaptic mitochondria isolated from 8 weeks old C57BL/6J female mice as described in [17]. Briefly, 50µg of mitochondrial protein was preincubated in the presence of test compounds for 3min in assay buffer containing 34mM potassium phosphate, 1.5mM KCN, 3.75µM rotenone, 1.5mM cytochrome C, 3mM phenazine methosulfate, pH7.2. The reaction was then started by the addition of 50µM palmitoyl-CoA, 100µM octanoyl-CoA or 200µM butyryl-CoA. Depending on the substrate used, reactions were monitored at 550 nm over a period up to 3h using a spectrophotometer (Synergy HTX Multi-Mode Reader, BioTek).

#### 3-hydroxyacyl-CoA dehydrogenase assay

The activity of 3-hydroxyacyl-CoA dehydrogenase was measured in isolated mitochondria from 8 weeks old C57BL/6J female mice [18]. Assay medium consisted of 0.1M potassium phosphate, 0.5M MOPS, 100µM DTT, 0.1% (w/v) TritonX-100, 150µM NADH, 5mM N-ethylmaleimide, pH6.16. 50µM of acetoacetyl-CoA was provided as substrate to the reaction. A concentration of 50ug of mitochondrial protein was used per well. Reaction was measured at 37°C by following the decrease in absorbance at 340nm using a spectrophotometer (Synergy HTX Multi-Mode Reader, BioTek).

#### Substrate profiling assay

To determine mitochondrial metabolic activity and substrate specificity, MitoPlates S-1 (#14105, Biolog) assays were performed according to the manufacturer’s instructions. Briefly, assay mix was added into all wells of a MitoPlate and the plate was incubated at 37°C for 1h to allow substrates to dissolve. Freshly isolated mitochondria were resuspended in 1×Biolog Mitochondrial Assay Solution (MAS) and dispensed into each well of a MitoPlate (40ug of mitochondrial protein per well). Color formation at 590nm was read kinetically for 2h using a spectrophotometer (Synergy HTX Multi-Mode Reader, BioTek) and the background was corrected for the blank sample. Results are presented as average rate per minute per ug of protein.

#### CPT1 activity in isolated brain mitochondria

CPT1 activity was measured using a colorimetric assay based on the literature [19]. Briefly, 15μg of mitochondrial protein was loaded onto 96-well plates in 200μl of reaction buffer containing 2mM DTNB, 116mM Tris–HCl [pH 8.0], 2.5mM EDTA, and 0.2% Triton-X 100. CPT1 inhibitor, Malonyl-CoA, or its vehicle control was also added to the reaction buffer at a final concentration of 40µM. The plates were incubated at room temperature for 20min to eliminate all pre-existing reactive thiol groups, and the reaction was initiated by adding 100µM palmitoyl-CoA as a CPT1 substrate and 1mM carnitine as a CPT1 cofactor respectively. After 10min incubation at 37°C, release of CoA-SH from palmitoyl-CoA was spectrophotometrically determined as a readout of CPT1 activity at 412nm using a spectrophotometer (Synergy HTX Multi-Mode Reader, BioTek) in kinetics mode with 20-s intervals for a total assay time of 1h.

#### Fatty acid uptake in isolated brain mitochondria

Fatty acid uptake capacity was measured in isolated brain mitochondria using a Fatty Acid Uptake probe (#UP07 – Dojindo) as per manufacturer’s instructions. Briefly, 8µg of mitochondrial protein was loaded onto a 96 well plate and the plate was centrifuged at 2000xg for 5min to seed mitochondria in the bottom of the wells. The plate was then incubated for 15min at 37°C, after which 0.2µl of the Fatty Acid Uptake probe were added to the wells (in a total volume of 100µl). The plate was incubated for another 15min at 37°C, after which 100µl of Quenching buffer was added per well. Fluorescence was measured (excitation at 485nm; emission at 535nm) using a spectrophotometer (Synergy HTX Multi-Mode Reader, BioTek).

#### Immunoblot analysis

For immunoblot analysis, equal amount of mitochondrial protein extracts (20μg/well) were separated by SDS-PAGE (Invitrogen) in MOPS-SDS running buffer and transferred onto 0.2µm nitrocellulose membranes for 1h at 30V. Membranes were then blocked with 5% milk in TBS-T (20mM Tris-HCl pH7.5, 150mM NaCl, 0.5% Tween-20) for 1h. Primary antibody incubations were performed overnight at 4°C. Washes were performed for 5min at RT in TBS-T. Incubation with secondary antibodies conjugated with horseradish peroxidase (HRP) were incubated for 2h at RT. Detection was done using the chemiluminescent ECL-Plus detection kit (Amersham) on a digital Amersham Imager 680 or 800 (GE Healthcare).

#### Mouse primary neuronal cultures

Primary neuronal cultures were obtained from whole brain of E18 embryonic C57BL/6 mice, as previously described [20]. Briefly, brains were collected in Hank’s Balanced Salt Solution supplemented with 1M HEPES (HBSS-HEPES). Meninges were removed and brain tissue was homogenized and dissociated with 0.25% trypsin + 10µl/ml DNaseI in HBSS-HEPES for 15min at 37°C. Tissue was washed with 10% Horse serum (HS) in HBSS-HEPES and centrifuged 1200xg for 10min at RT. Supernatant was discarded and pellet was washed twice with HBSS-HEPES. Pellet was resuspended in 10% HS in Minimum Essential Medium (MEM) with 0.6% Glucose and 100U/ml Penicillin-Streptomycin (MEM-HS) and strained using a 70µm cell strainer. Cell viability was assessed using Trypan blue. Cells were plated at different densities depending on the assay, on plates coated with 0.01mg/ml Poly-D-Lysine hydrobromide. After 4h, medium was replaced with Neurobasal medium supplemented 0.5mM L-Glutamine, 20U/ml Penicillin-Streptomycin and 2% B-27 (NB-B27).

#### Mitochondrial respiration measurements

Mitochondrial respiration was assessed by measuring the oxygen consumption rate (OCR) using the XFe24 Extracellular Flux Analyzer (Seahorse Bioscience, Agilent, USA). For brain mitochondria, freshly collected synaptic and non-synaptic mitochondrial fraction (8μg of total protein concentration) were resuspended in mitochondrial assay solution (MAS)[21] and plated in a 24-well Seahorse plate followed by centrifugation 2000 x g, 5min, 4°C. After centrifugation, 450μl MAS supplemented with 10mM malate, 10mM succinate and 10mM pyruvate, pH 7.4 was added to the plate. Fatty acid substrates 40µM palmitoyl-CoA + L-carnitine, 100µM octanoyl-CoA and 200µM butyryl-CoA were diluted in the supplemented MAS. OCR measurements were performed over 2min after a 30s mix and 10s periods. Three measurements were collected for basal respiration, followed by two measurements after addition of 2mM ADP, followed by two measurements after addition of 20μM Oligomycin, followed by two measurements after addition of 20μM Carbonyl cyanide-4-(trifluoromethoxy) phenylhydrazone (FCCP), followed by two measurements after addition of 20μM Rotenone and 20μM Antimycin A.

For primary neuronal cultures, OCR experiments were performed on E18 mouse neurons at 10 days in vitro (DIV), and an Agilent Seahorse XF Palmitate-BSA FAO Substrate protocol was performed with slight modifications. Neurons were placed in FAO assay buffer containing 111mM NaCl, 4.7mM KCL, 1.25Mm CaCl2, 2mM MgSO4 and 1.2mM NaH2PO4, supplemented with 2.5mM Glucose, 0.5mM L-Carnitine and 5mM HEPES, pH=7.4, and incubated for 4h at 37°C. FAO buffer was also supplemented with long- and medium-chain fatty acids in the form of 25µM palmitate and 25µM octanoate or BSA vehicle control. To assess the effect of inhibiting the metabolism of long-chain fatty acids, neurons were pre-incubated with 10µM etomoxir 15 minutes prior to the addition of fatty acids to the medium.

OCR measurements were performed over 3min after a 3min mix and 2min wait periods. Four measurements were collected for basal respiration, followed by three measurements after addition of 1.25μM Oligomycin, three measurements after addition of 0.5μM FCCP, and three measurements after addition of 1μM Antimycin A and 1μM Rotenone.

#### ATP content in neurons

The ATP content was measured in DIV10 neurons treated for 4h with 25µM Palmitate or Octanoate. The effect of 10µM etomoxir was also assessed. ATP content was determined using the Luminescent ATP Detection Assay Kit as per manufacturer’s instructions (Abcam, ab113849). Cells were lysed and collected with 350μl 1x Detergent Mix, mixed for 5 min at 600rpm at RT, and centrifuged at 10000xg for 1min at RT. 150μl of each sample, as well as 10μl of each standard and 140μl of 1x Detergent Mix were pipetted in a flat 96-well plate and incubated with 50μl of Substrate Solution at RT protected from light for 5min with agitation, followed by 10min without agitation. Luminescence of samples, blanks and standard wells were measured using a spectrophotometer (Synergy HTX Multi-Mode Reader, BioTek). From the ATP content standards, a standard curve was plotted and then used to determine the ATP content of each sample. All ATP content levels were normalized to protein levels quantified using the Bradford Protein Quantification Assay.

#### BODIPY 558/568 C12 uptake in neurons

To assess fatty acid uptake, primary neurons were plated at 1x105 cells/well on 13mm coverslips (in a 24 well tissue culture plate). DIV10 neurons were incubated with 2μM BODIPY 558/568 C12 (referred as Red-C12 from here onwards) for 24h in NB-B27 media. Neurons were then transferred to NB-B27 media without Red-C12 for 1h. Neurons were washed and chased for 4 hours in NB-B27 media. To evaluate fatty acid uptake in neuronal periphery, 8x104 cells were plated in microfluidic devices [22], [23], 2-compartment devices with 450µm microgroove barrier. Fresh NB-B27 was added every 3 days. A volume difference between the somal (170-180µL) and the axonal (120-130µL) compartments was maintained to create a hydrostatic pressure, thus fluidically isolating each compartment. The Red-C12 incubation procedure described above was applied to neurons, exclusively in the axonal compartment. Neurons were then subjected to an immunofluorescence protocol as described below.

#### Immunofluorescence assay

Neurons (DIV10) were washed in PBS+/+ (0.33mM MgCl2.6H2O, 0.9mM CaCl2.2H2O) and fixed in 4% paraformaldehyde (PFA) in PBS+/+ for 20min at RT. Neurons were permeabilized with 0.5% Triton X-100 in PBS+/+ for 10min at RT and washed with PBS. Blocking was performed for 1h at RT with blocking buffer (0.2% gelatine, 2% FBS, 2% BSA, 0.3% Triton X-100 in PBS) supplemented with 5% goat serum followed by primary antibody incubation overnight at 4°C. Incubation with secondary antibodies conjugated with Alexa-Fluorophores and DAPI (1µg/mL) was performed for 1h at RT. Coverslips were mounted on Vectashield (#H1000 - Vector Laboratories).

For microfluidics, neurons were pre-fixed for 5min at 37°C with 5% CO2, by adding 4% PFA to each compartment (together with NB-B27). Pre-fixation solution was removed, and 4% PFA was added to each compartment for 20min at RT. Each compartment was washed with PBS+/+. Microfluidic chamber was carefully removed, and coverslip was washed with PBS+/+. Neurons were permeabilized with 0.5% Triton X-100 and immunofluorescence protocol proceeded as mentioned above.

#### Whole-cell patch-clamp recordings

For whole-cell patch-clamp recordings, primary neurons were seed in 13mm coverslips that were mounted in a Carl Zeiss Axioskop 2FS upright microscope (Jena, Germany) equipped with an AxioCam MRm (Zeiss), a 40x immersion objective with 2 and 4 zoom (i.e. up to 160x magnification) and a differential interference contrast-infrared (DIC-IR) CCD video camera (VX44, Till Photonics, Gräfelfing, Germany) [41]. Whole-cell patch-clamp recordings were performed using electrodes pulled from borosilicate glass capillaries (1.5mm outer diameter, 0.86mm inner diameter, GC150F-10, Harvard Apparatus, Holliston, MA, USA) in a PC-10 vertical (Narishige Group, London, UK) microelectrode puller. The intracellular solution used in this study contained 125mM K-Gluconate, 11mM KCl, 10mM HEPES, 10 mM Phosphocreatine, 2mM MgATP, MgCl2 2mM, 2mM EGTA, 0,3mM NaGTP and 0,1mM CaCl_2,_ pH=7.3 (adjusted with KOH 1 M), 280–290 mOsm. Patch-clamp recordings were performed in a submerged recording chamber that was continuously superfused by an open gravitational superfusion system at 2-3 mL/min with artificial cerebrospinal fluid (aCSF – 124mM NaCl, 26mM CaHCO_3_, 10mM Glucose Monohydrate, 10mM Phosphocreatine, 3mM KCl, 2mM CaCl_2_, 1.25mM NaH_2_PO_4,_ 1mM MgSO_4_) oxygenated with 95% O_2_ and 5% CO_2_, at room temperature. Data was recorded with an Axopatch 200B (Axon Instruments) amplifier. For all patch-clamp recordings, whole-cell access was established following formation of a gigaseal (>1 GΩ) between pipette tip and cell membrane. Immediately after having whole-cell access, the membrane potential of the neurons was measured in current clamp mode (VH approximately = -60 mV). The acquired signals were filtered using a 2 kHz lowpass Bessel filter, and data were digitized at 10 kHz using Digidata 1440A and registered by the Clampex software (version 11.0.3, Molecular Devices).

The spontaneous postsynaptic currents were recorded in the voltage-clamp mode for 5 min.

### Quantification and statistical analysis

For all the quantitative analysis, at least 3 independent experiments from 3 different primary neuronal cultures, or 3 different mitochondrial isolation protocols were performed. The statistical analysis was implemented using Prism 8.4.3 software (GraphPad). An appropriate statistical test was chosen based on the dataset. Detailed information is provided in each figure legend.

